# STAT3 Sustains Tumorigenicity Following Mutant KRAS Ablation

**DOI:** 10.1101/2025.02.21.639533

**Authors:** Stephen D’Amico, Varvara Kirillov, Jingxuan Liu, Zhijuan Qiu, Xinyuan Lei, Hong Qin, Brian S. Sheridan, Nancy C. Reich

## Abstract

Oncogenic KRAS mutations underlie some of the deadliest human cancers. Genetic or pharmacological inactivation of mutant KRAS is not sufficient for long-term control of advanced tumors. Using a conceptual framework of pancreatic ductal adenocarcinoma, we find that CRISPR-mediated ablation of mutant KRAS can terminate tumor progression contingent on the concomitant inactivation of STAT3. STAT3 inactivation is needed to ensure that KRAS-ablated tumor cells lose their malignant identity. Mechanistically, the combined loss of mutant KRAS and STAT3 disrupts a core transcriptional program of cancer cells critical to oncogenic competence. This in turn impairs tumor growth in mice and enhances immune rejection, leading to tumor clearance. We propose that the STAT3 transcriptional program operating in cancer cells enforces their malignant identity, rather than providing classical features of transformation, and shapes cancer persistence following KRAS inactivation. Our findings establish STAT3 as a critical enforcer of oncogenic identity in KRAS-ablated tumors, revealing a key vulnerability that could be exploited for combination therapies.

**Significance:** The limited clinical success of KRAS inhibitors points to the need to identify means by which tumor cells maintain stemness and immune evasion. We make an unprecedented finding that the STAT3 transcription factor can sustain tumorigenicity of pancreatic cancer cells following depletion of the KRAS oncogenic driver. The results have important implications for successful therapeutic intervention.

## Introduction

Pancreatic ductal adenocarcinoma (PDAC) is a highly aggressive malignancy characterized by rapid progression and exceptional resistance to anticancer treatment (Halbrook et al, 2023). Despite KRAS mutations driving 90% of PDAC cases, direct KRAS inhibition has had limited success, in part due to tumor heterogeneity and adaptive resistance (Briere et al, 2021; Liu et al, 2024; Punekar et al, 2022). Historically, therapies have assumed that KRAS-driven cancers are entirely dependent on KRAS signaling. However, the degree of oncogenic KRAS addiction can vary across individual tumor cells, challenging this notion (Muzumdar et al, 2017; Punekar et al., 2022; Salmon et al, 2023; Singh et al, 2009; Yuan et al, 2018). Results suggest that even the most potent inhibition of oncogenic KRAS alone may not be sufficient for curing cancer in humans. We investigate signal transducer and activator of transcription (STAT3) as a key regulator sustaining malignancy following mutant KRAS inactivation (D’Amico *et al*, 2024; Fukuda *et al*, 2011; Huynh *et al*, 2019; Lee *et al*, 2014; Lesina *et al*, 2011; Philips *et al*, 2022; Salmon *et al*., 2023).

The current study aimed to provide a more direct investigation of the relationship between STAT3 and oncogene-driven tumorigenesis. Pancreatic tumorigenesis is a process that involves the gradual accumulation of mutations in oncogenes (e.g. KRAS) and tumor suppressor genes (e.g. TP53, CDKN2A, and SMAD4)(Hayashi *et al*, 2021). Cooperative mutations of oncogenes and tumor suppressors appear to function within a specific cellular context termed oncogenic competence (Baggiolini *et al*, 2021; Hsieh *et al*, 2024; Weeden *et al*, 2023). The role of the STAT3 transcription factor in the oncogenic process remains elusive with reports indicating either tumor promotion or suppression depending on the tissue or cell type studied. Although STAT3 is not inherently oncogenic, it has been reported to facilitate PDAC tumor progression following disruption of TGF-beta/SMAD4 signaling (Laklai *et al*, 2016; Singh & Settleman, 2009). In addition, transcriptional signatures of STAT3 and KRAS activity in pancreatic cancer cells partially overlap (D’Amico *et al*., 2024). We investigated the hypothesis that STAT3 promotes maintenance and self-renewal of pancreatic cancer cells following mutant KRAS inhibition. Results show that CRISPR-mediated KRAS ablation impedes tumor growth, and complete inhibition of resistant tumor cells is achieved with concomitant inactivation of STAT3. Mechanistically, the combined loss of KRAS and STAT3 disrupts a core transcriptional program resulting in impaired tumor growth in mice and enhanced immune rejection. Our findings identify a specific role for STAT3 in tumor cell maintenance that becomes apparent with the inhibition of mutant KRAS and provide a rationale for developing orthogonal cancer therapies targeting the function of mutant KRAS and STAT3.

## Results

### STAT3 correlates with KRAS independence in human PDAC

Studies with epithelial cancer cells expressing mutant KRAS have shown that at an advanced tumor stage, cancer cells can have a reduced dependency on the KRAS oncogene (Lim & Counter, 2005; Muzumdar *et al*., 2017; Singh *et al*., 2009; Yuan *et al*., 2018). Specific gene expression signatures have been derived for cancer cells designated KRAS-dependent (KRAS_sig/KRAS-type) or KRAS-independent (RSK_sig/RSK-type) (Singh & Settleman, 2009; Yuan *et al*., 2018). To examine the potential association between STAT3 expression and the degree of oncogenic KRAS dependency, we analyzed gene expression profiles in datasets of human PDAC tumor samples. Two clear trends emerged. First, tumor samples stratified into either KRAS dependent or KRAS-independent subtypes regardless of cancer stage or treatment history (Fig. 1A, Fig. S1A, S1B)(Singh *et al*., 2009; Yuan *et al*., 2018). Second, STAT3 mRNA and STAT3 regulated gene expression correlated with tumor samples classified as KRAS-independent (Fig. 1A, Figs. S1A, S1C, S1D)(Dauer *et al*, 2005). We also detected heterogeneity in human pancreatic tumor specimens from more than 20 patients with primary and metastatic PDAC. Classically, ERK mitogen-activated protein kinase activity is a downstream effector of KRAS and identifies KRAS-dependent cancer cells (Klomp *et al*, 2024). Using immunohistochemistry for activated ERK we found extensive heterogeneity, consistent with evolution of KRAS independence (Figs. S1E, S1F).

**Figure 1.**
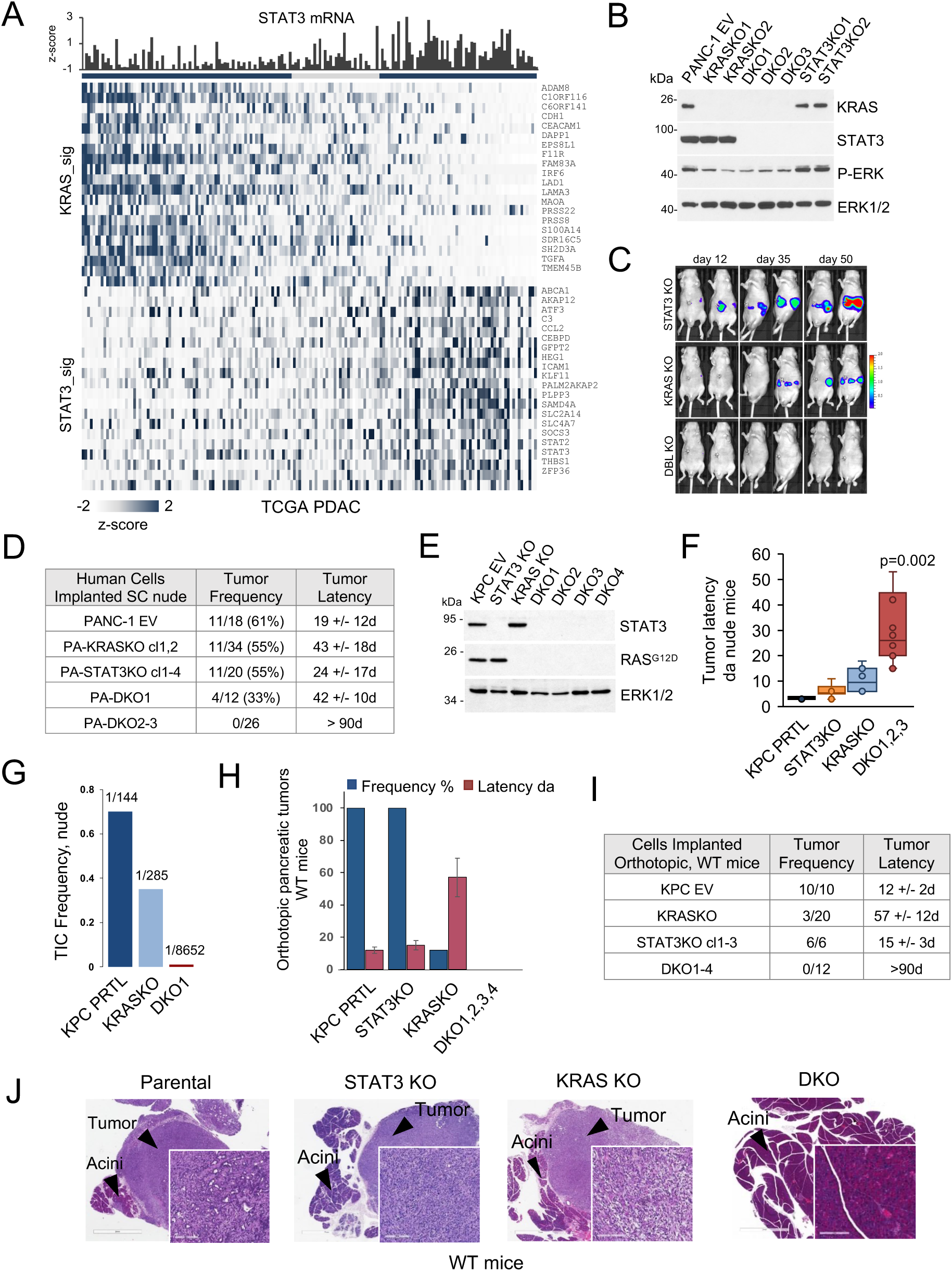
STAT3 maintains survival of mutant KRAS-ablated pancreatic tumors. A. Gene expression profiles of human PDAC tumor samples from the TCGA database (n=150) segregated according to KRAS dependency (KRAS_sig)(Singh & Settleman, 2009) and STAT3 dependent gene regulation (Dauer *et al*., 2005). Top panel displays corresponding STAT3 mRNA expression (z-scores) of each tumor. B. Western blot of human PANC-1 parental cells expressing empty lentiviral vector (EV) and cells with CRISPR-mediated knockout of endogenous KRAS^G12D^ (KRAS KO, two clones), STAT3 (STAT3 KO, two clones), or KRAS^G12D^ and STAT3 (DKO, three clones). ERK1/2 and phospho-ERK are loading controls. C. Representative *in vivo* bioluminescence imaging of nude mice following orthotopic implantation of STAT3 KO cells, KRAS^G12D^ KO cells, or DKO cells derived from human PANC-1 cell line transduced with the luciferase gene. Days denote time following implantation. D. Tumor formation of human PANC-1 EV cells and derived independent clones (cl) with KRAS KO, STAT3 KO, or KRAS^G12D^ and three STAT3 DKO. Tumor frequency and latency are noted. E. Western blot of KRAS^G12D/+^ p53^R172H/+^ KPC cells with parental (EV) and derived isogenic clones showing ablation of endogenous KRAS^G12D^ (KRAS KO), STAT3 (STAT3 KO), and four independent clones with a double ablation of KRAS^G12D^ and STAT3 (DKO). ERK1/2 is a loading control. F. Box plots showing subcutaneous tumor latencies (days, ∼1mm) in nude mice of parental KPC cells or isogenic knockout derivatives noted. G. Limiting dilution assays in nude mice were used to calculate frequency of tumor initiating cells (TICs) of parental KPC, KRAS KO, and DKO1 cells. H. Parental KPC cells or knockout derivative cells were orthotopically implanted into the pancreas of syngeneic wild type mice (C57BL/6). Tumor formation frequency (%) and latency in days (1 mm^3^) are shown on the same axis (n=6-20 for each). I. Specifics of tumor development in wild-type mice following orthotopic implantation into the pancreas with KPC empty vector, STAT3 KO, KRAS KO, or DKO cells shown in (H). Tumor frequency shown as number of mice with tumors vs. total number of mice. Tumor latency shown in days (d). J. Representative histological images of H&E staining of KPC pancreatic tumors following orthotopic implants of designated cells in wild-type mice. Inset scale bar 200mm.

### Loss of STAT3 prevents tumor formation after KRAS ablation

To evaluate the contribution of STAT3 to the tumorigenicity of PDAC cells, we used CRISPR-mediated gene editing to eliminate endogenous KRAS and/or STAT3 in the human PANC-1 cell line. Although PANC-1 cells bear an activated KRAS G12D mutation, they have been classified as KRAS-independent type (Singh & Settleman, 2009). We first isolated KRAS KO and STAT3 KO PANC-1 independent clones, confirmed loss of KRAS and STAT3 by protein expression, and tested their ability to form orthotopic tumors in nude mice (Figs. 1B, 1C, S1G). Both the STAT3 KO and the KRAS KO cells formed tumors, but tumor latency increased with KRAS ablation. To evaluate the dependency of the KRAS KO PANC-1 cells on STAT3 to form tumors, we depleted STAT3 in the KRAS KO cells that were tumorigenic and generated KRAS/STAT3 double knockout (DKO) clones. While STAT3 loss did not alter PANC-1 cell proliferation in vitro (Fig. S1H), its absence in KRAS-ablated cells prevented orthotopic tumor formation *in vivo*. This suggests STAT3 is essential for maintaining tumorigenicity despite KRAS depletion. We also tested the ability of these cells to develop tumors as subcutaneous xenografts (Fig. 1D). KRAS KO PANC-1 cells retained the ability to form subcutaneous tumors in nude mice, but more slowly than parental PANC-1 controls, consistent with published observations (Muzumdar *et al*., 2017). However, only one of the three DKO PANC-1 clones depleted for both KRAS and STAT3 was able to form any tumors. The results suggest that under conditions of KRAS depletion, inactivation of STAT3 can completely abolish pancreatic tumor growth.

To gain a better understanding of the roles of STAT3 in tumor maintenance, we generated a panel of CRISPR-edited murine KPC cell lines (KRAS^G12D/+^; p53^R172H/+^) (Hingorani *et al*, 2005), each mimicking complete inactivation of the KRAS and/or STAT3 genes (D’Amico *et al*., 2024; D’Amico *et al*, 2018; Ischenko *et al*, 2021). Three types of isogenic cell lines were examined: KRAS KO, STAT3 KO, and KRAS/STAT3 double knockout (DKO). DNA sequencing revealed insertion-deletion mutations within the targeted STAT3 and KRAS alleles, and Western blot analysis showed no detectable STAT3 or KRAS^G12D^ protein expression (Fig. 1E, Fig. S1I). Loss of STAT3 alone minimally affected the growth of cells in culture or as tumors in nude mice (Fig. 1F, Fig. S1J) (D’Amico *et al*., 2024; D’Amico *et al*., 2018). KRAS KO clones produced tumors in nude mice when implanted subcutaneously or orthotopically into the pancreas, although with a longer latency than the parental controls (Ischenko *et al*., 2021; Muzumdar *et al*., 2017). Thus, ablation of KRAS alone did not eliminate their tumorigenic properties in nude mice. In contrast, the tumorigenicity of DKO cell lines independently derived from the KRAS KO tumorigenic cells was either attenuated or abolished (i.e., no tumor formation after 60 days of observation) (Fig. 1F, Fig. S1K). We used limiting dilution assays in nude mice to estimate the frequency of tumor-initiating cells (TICs)(Hu & Smyth, 2009). The TICs ranged from ∼0.7% in parental KPC cells to ∼0.034% in DKO1 cells (∼ 20-fold change) requiring more than 2×10^4^ DKO implanted cells for tumor formation (Fig. 1G).

We next determined the impact of an intact immune system on pancreatic tumor growth of the isogenic modified KPC cells. Tumor formation was assessed following orthotopic implantation of KPC cell lines into the pancreas of wildtype C57Bl/6 mice. The KRAS KO cells were severely impaired in tumor forming capacity, but their tumorigenicity was not eliminated completely, as shown previously (Figs. 1H, 1I)(Ischenko *et al*., 2021). However, co-deletion of STAT3 in the KRAS KO tumorigenic cells fully abrogated tumor growth in wild-type recipient mice. Histology showed that pancreatic tumors derived from KPC parental cells resembled well-differentiated PDAC with prominent glandular structures, whereas STAT3 KO and KRAS KO tumors exhibited a mesenchymal-like morphology as reported previously (Fig. 1J)(D’Amico *et al*., 2024). Importantly, there was no evidence of malignancy in the pancreata of wild-type mice transplanted with DKO cell lines after 90 days of observation. The data indicate that STAT3 is needed to sustain pancreatic tumor growth under conditions of intact immunity following the genetic inactivation of mutant KRAS.

### Combined loss of KRAS and STAT3 disrupts a core transcriptional program in PDAC cells

To gain insight into the mechanisms by which STAT3 mediates continued tumor growth following KRAS inactivation, we performed RNA sequencing analysis of KPC cells and their STAT3 KO, KRAS KO, and DKO derivatives. Using fold change ≥4 and false discovery rate <0.05 cut-offs, we identified 2000 genes as differentially expressed by DKO vs. parental cells. Pairwise differential expression analyses are displayed as comparative scatter plots (Fig. 2A). Pathway enrichment analysis of up- and down-regulated genes revealed that the expression of genes involved in stem cell maintenance, epithelium development, and cell adhesion was reduced in DKO samples (GO, Reactome, MSigDB) (Fig. 2B, Figs. S2A, S2B). The DKO cells maintained significant levels of phosphorylated MEK1/2 and ERK1/2 (Fig. S2C). The most salient feature underlying the changes in DKO cells was the perturbation of transcription factors (at least 60 genes) controlling lineage specificity, differentiation, proliferation, and survival (Figs. 2C, 2D, Fig. S2D). These were examined based on a predictive list of core transcription factors that regulate tumor cell fates (Reddy *et al*, 2021). Analyses showed that a majority of deregulated transcription factors (e.g. ELF3, FOXA1/2, HNF4A, KLF4, KLF5, SNAI, etc.) were those involved in cell fate decisions, and that DKO cells underwent reprogramming to a far greater extent than either STAT3 or KRAS single knockouts. Western blot analyses confirmed the reduced expression of HNF4A, FOXA1, FOXA2, KLF4, GATA5 in DKO cells, transcription factors involved in differentiation and lineage specification (Fig. 2E, Fig. S2D). These findings indicate that STAT3 works to sustain a core transcriptome profile within KRAS-ablated PDAC tumor cells.

**Figure 2.**
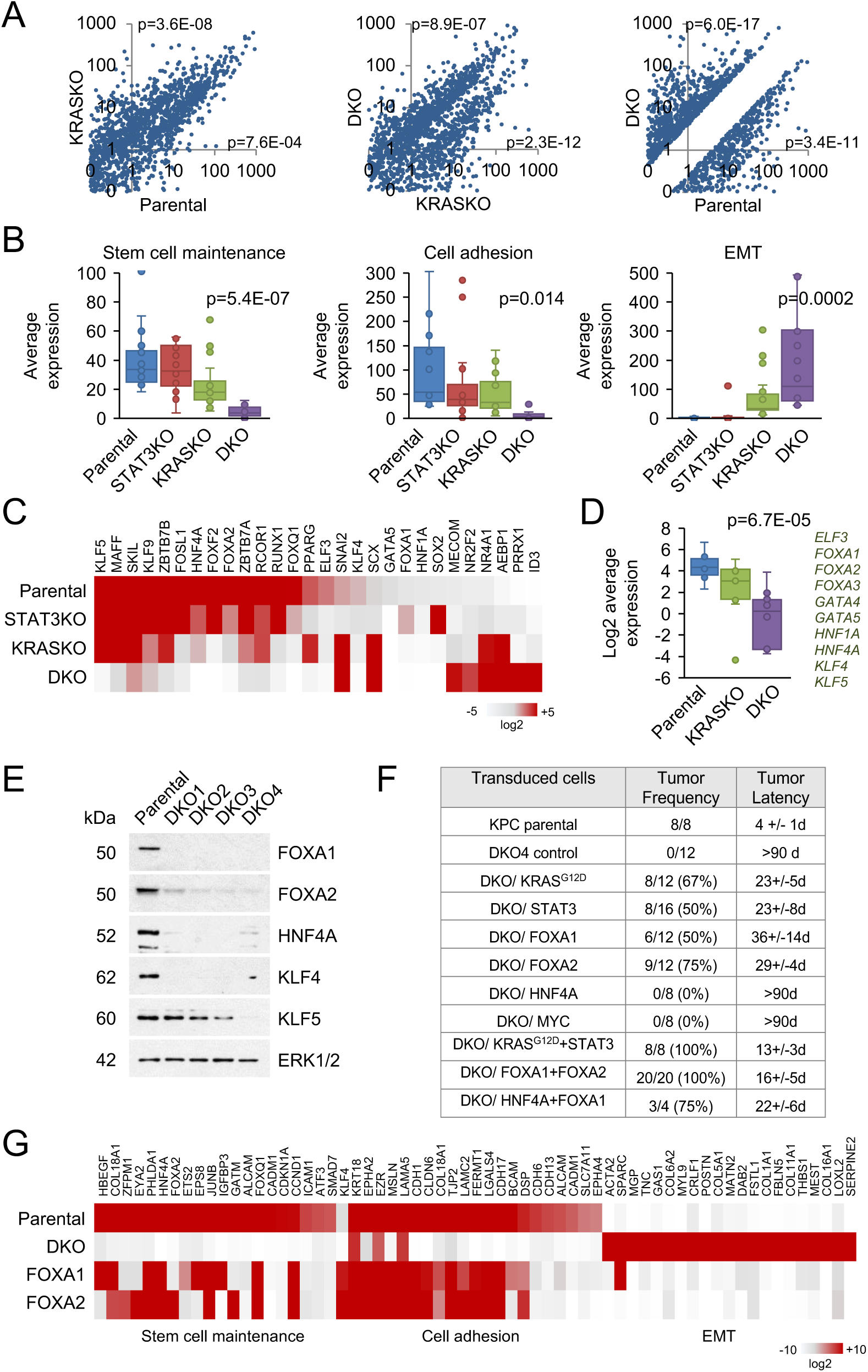
Transcriptional profiling of cells with combined loss of KRAS and STAT3. A. Differential expression of genes displayed as scatter plots for KPC parental, KRAS KO and DKO cells derived from RNA-seq data with >4-fold difference and FDR <0.05. B. Pathway scoring using the top 20 genes corresponding to stem cell maintenance, cell adhesion, and EMT signatures from The Molecular Signatures Database (MSigDB) in KPC parental cells and derived KRAS KO, STAT3 KO, and DKO cells. C. Comparative heatmaps of RNA-seq expression data of transcription factors in KPC parental and derived KRAS KO, STAT3 KO, and DKO4 cells. D. Summary boxplots of differential expression for a subset of transcription factors that regulate lineage specificity from RNA-seq data of KPC parental, KRAS KO and DKO cells. E. Western blot showing the protein expression of a subset of transcription factors in KPC parental cells and four double KO (DKO) clones. ERK1/2 is a loading control. F. Restoration of tumorigenicity in nude mice of DKO4 cells following transduction with mutant KRAS^G12D^, STAT3, HNF4A, MYC, FOXA1, FOXA2 or the indicated combinations compared with parental KPC cells. Tumor frequency (tumors/implantation site) and latency (days) are provided. G. Heatmaps of RNA-seq expression data of KPC parental cells, DKO4 cells, and DKO4 cells reconstituted with FOXA1 or FOXA2. Specific gene signatures were obtained from MSigDB as in Fig. 3B.

We ectopically re-expressed STAT3, KRAS^G12D^, and select transcription factors in the DKO cells to determine whether reconstitution could reverse cell morphology and recover tumorigenic capacity. Expression of STAT3 or mutant KRAS by lentiviral transduction in DKO4 cells partially restored tumorigenicity in nude mice, and their pairwise expression further restored tumor formation (Fig. 2F, Figs. S2E, S2F). Among the transcription factors tested, expression of FOXA1 and FOXA2 achieved the highest degree of phenotypic conversion. In contrast, no phenotypic reversal was observed following ectopic expression of HNF1A, HNF4A, KLF5, or MYC. These data demonstrate that the combined loss of KRAS and STAT3 (or their downstream targets FOXA1/2) significantly impacts tumorigenic growth *in vivo*. Ectopic expression of FOXA1 or FOXA2 also generated tumors that were morphologically similar to parental KPC tumors in terms of epithelial differentiation (Fig. S2G). Further examination showed that ectopic expression of FOXA1 and FOXA2 caused DKO cells to revert to a state that resembled parental KPC cells in terms of stem cell maintenance, cell adhesion, and epithelial differentiation gene expression supporting their respective roles in the maintenance of the KRAS mutant phenotype (Fig 2G). The DKO cells supplemented with FOXA1/2 also expressed genes more closely associated with PDAC ductal 2 subtype, distinguishing a more malignant cell state (Fig. S2H)(Cui Zhou *et al*, 2022; Peng *et al*, 2019). To ascertain whether transcriptome results of FOXA1 and FOXA2 reconstituted cell lines could be projected onto human PDAC, we computed gene expression signatures derived from the top 100 up- or down-regulated genes. PDAC tumor samples from TCGA were segregated according to RAS dependency signature and differentiation status. Alignment revealed that FOXA1/2 regulated expression signatures co-segregated with epithelial (EPI) KRAS-dependent (RDI) tumors (Pearson’s r>0.6) and were inversely correlated with mesenchymal (MES) KRAS independent (RSK) tumors (Pearson’s r>0.6) (Fig. S2I). Transcriptomic analysis of human PANC-1 cells and their derivatives identified more than 2,600 differentially expressed genes in the DKO cells at a threshold of 4-fold difference or greater. Among the down-regulated cellular pathways in DKO cells were oxidative phosphorylation, ribosome biogenesis, and cytoplasmic translation, whereas the upregulated pathways included axon guidance and muscle tissue development (Fig. S2J). A subset of transcription factors that regulate various biological processes are differentially expressed in the PANC-1 DKO cells and not evident in single knockouts (Fig. S2K). Together, the data demonstrate that combined loss of KRAS and STAT3 (or their downstream targets) leads to cellular reprogramming that significantly impacts tumorigenic growth.

### STAT3 deletion enhances tumor rejection

We investigated whether depletion of STAT3 could improve the efficacy of KRAS inhibition, since activation of STAT3 was reported in response to tumor regression with RAS/RAF/MAPK inhibitors (Lee *et al*., 2014; Salmon *et al*., 2023). We first evaluated the degree of STAT3 activation and immune infiltration of KPC tumors from mice treated with a protocol that encompasses the KRAS^G12D^ inhibitor MRTX1133 and MEK inhibitor GSK1120212 (Trametinib) with immune checkpoint inhibitors (ICIs) (anti-PDCD1, CTLA4, and CD40 antibodies). We previously observed this treatment protocol to cause regression of PDAC tumors in mice at 60-80% efficiency within two weeks (Li *et al*, 2023). Mice treated with MRTX1133, GSK1120212, or ICIs alone did not significantly reduce tumor size. Immunohistochemistry was performed to monitor STAT3 activation by tyrosine phosphorylation and immune infiltration in tumors of mice untreated or treated with this protocol. As expected, treatment of mice bearing KPC tumors induced robust STAT3 phosphorylation (>100 fold compared with untreated controls) and CD8 T cell infiltration (>20 fold) (Li *et al*, 2018)(Fig. 3A). We next investigated whether inactivation of STAT3 could improve the efficacy of tumor regression with the KRAS pathway inhibitors administered in combination with immune checkpoint inhibitors. Mice bearing orthotopic tumors formed by parental KPC cells expressing empty vector or isogenic STAT3 KO cells were either untreated (d1) or received four treatments with the KRAS inhibitory cocktail during a seven day period and were evaluated by tumor weight on the seventh day (d7T) (Fig. 3B). The results are representative of two experiments that used different treatment regimens. Mice bearing the STAT3 KO tumors showed a significant reduction in tumor size compared to mice with parental KPC empty vector tumors, even complete regression, within one week (Fig. 3C). The results provide preclinical evidence that STAT3 depletion can enhance the antitumor effects of KRAS combination therapy.

**Figure 3.**
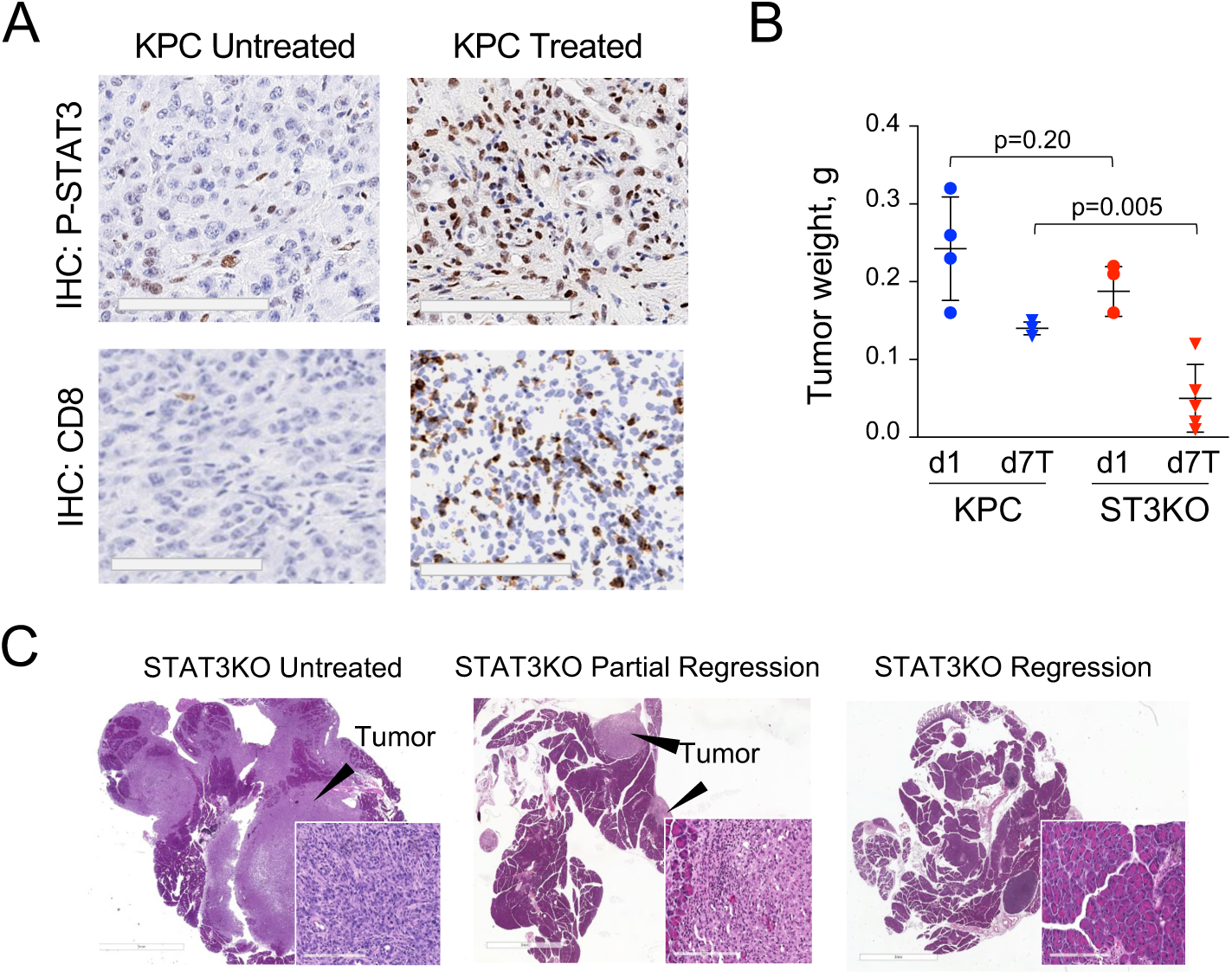
Tumor regression following combination therapy. A. Representative immunohistochemistry (IHC) of orthotopic tumors formed by KPC parental cells (Li *et al*., 2018) in untreated mice or mice treated by administration of a cocktail of MRTX1133, GSK1120212, and antibodies to PDCD1, CTLA4, and CD40 as previously described (Li *et al*., 2023). Tumors were stained by IHC with antibodies to tyrosine 705 phosphorylated-STAT3 or CD8a. Scale bar 200mm. B. Effect of combination drug treatment on growth of orthotopic tumors formed by KPC cells or isogenic STAT3 KO cells. Tumor weights are shown for control mice at the onset of the experiment day 1 (d1) and for mice harvested on day 7 following drug administration on days 1, 2, 4, and 6 (d7T). Mice that were not treated formed tumors from KPC or STAT3 KO cells at day 7 equal to or greater than 0.5g (not shown). Each data point represents one mouse, n=4-5 per group. C. Representative histological H&E images of tumors formed by STAT3 KO cells in untreated mice or mice following drug administration.

### Loss of KRAS and STAT3 in PDAC cells modulates tumor immunity

PDAC cells depleted for both mutant KRAS and STAT3 (DKO) do not form tumors in isogenic wild-type mice, and therefore tumor analysis is not possible (Fig. 1). We therefore investigated an independent model system with a doxycycline-inducible KRAS^G12D^ and mutant p53^R172H^ (termed iKRAS) (Collins *et al*, 2012b). The drawback of this model is that it does not reflect tumor evolution with reduced dependency on KRAS. We used CRISPR-mediated gene editing to isolate several independent iKRAS clones depleted for STAT3 expression (Fig. 4A). Both iKRAS and derived STAT3 KO cell clones formed tumors in nude mice following administration of doxycycline. However, when transplanted into the pancreas of syngeneic wild-type mice, there was decreased and varied tumorigenicity of the STAT3 KO clones (Fig. S3A). To investigate the nature of orthotopic tumors formed in wild type mice, we evaluated immune cell infiltration of tumors formed by iKRAS control cells and the most fit STAT3 KO clone by flow cytometry. Results representing two experimental analyses with tumors pooled from individual mice treated with doxycycline are shown in Fig. 4B. Tumors formed by the STAT3 KO cells showed an increase in T cell number with a higher CD8 to CD4 ratio compared with iKRAS parental cells expressing empty vector. Since withdrawal of doxycycline extinguishes expression of the KRAS transgene, we assessed the effect of STAT3 loss on short-term regression of tumors (Collins *et al*., 2012b). As early as four days post-doxycycline withdrawal, the median weight of STAT3 KO tumors was reduced compared to controls (Fig. 4C). *In vivo* imaging with luciferase-expressing cells showed some of the STAT3 KO tumors became undetectable by seven days of doxycycline withdrawal (Fig. S3B). Overall, the data indicate that co-deletion of STAT3 and KRAS impairs development of tumors and facilitates tumor rejection in wild-type recipients.

**Figure 4.**
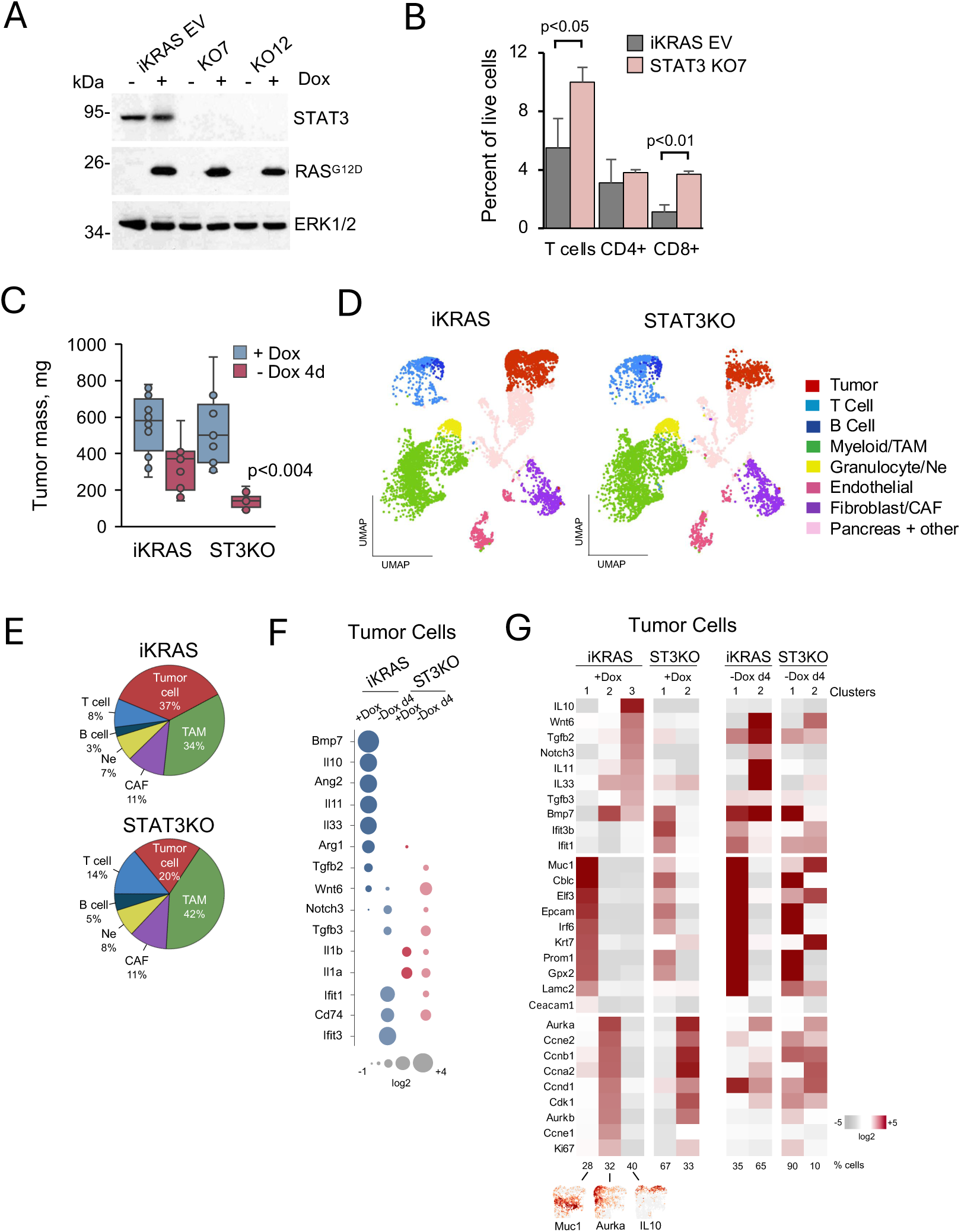
Influence of STAT3 loss in an inducible mutant KRAS PDAC model. A. Western blot of iKRAS PDAC parental cells expressing empty vector (EV) and two independent STAT3 KO derivatives (KO7 and KO12) untreated (-) or treated (+) with doxycycline. Protein expression detected with antibodies to STAT3 and KRAS^G12D^. Antibodies to ERK1/2 used as loading controls. B. Flow cytometry of tumors formed by orthotopic implantation of parental iKRAS cells or derived STAT3 KO7 cells into the pancreas of wildtype FVB mice. Mice were administered doxycycline and pancreatic tumors that formed were pooled from 2-3 individual mice. Single cells were dispersed and stained with fluorescently tagged anti-TCRβ, anti-CD3ε, anti-CD4, or anti-CD8a. Representative results from one experiment are shown. C. Pancreatic tumor weights (mg) are shown following orthotopic implantation of iKRAS EV parental or STAT3 KO cells into wildtype FVB mice. Mice were either treated with doxycycline (+Dox) or treated and withdrawn from doxycycline for 4 days (-Dox 4d) (n>6). D. UMAP projection images derived with Loupe browser software from scRNA-seq of pancreatic tumors. Analyses were performed following orthotopic implantation of iKRAS parental cells (left) or STAT3 KO7 cells (right) into the pancreas of wildtype FVB mice followed by treatment of mice with doxycycline. Main cell types are displayed in clusters distinguished by color. Tumors pooled from 2-3 individual mice. E. Pie charts displaying the relative percent of malignant tumor cells, TAMs, CAFs, Neutrophils (Ne), B cells and T cells and in the pancreatic tumors formed by iKRAS or STAT3 KO cells in FVB mice treated with doxycycline derived from scRNA-seq in Fig. 4D. F. Dot heatmap expression comparison of four scRNA-seq aggregated datasets. Expression of a subset of immunoregulatory genes in malignant cells from orthotopic tumors formed by iKRAS control cells or derived STAT3 KO cells in wildtype FVB mice. Mice were treated with doxycycline (+Dox) or treated and withdrawn from doxycycline for 4 days (-Dox 4d). G. Heatmaps of differential gene expression from scRNA-seq individual datasets of malignant cells from iKRAS or STAT3 KO orthotopic tumors formed in mice treated with doxycycline (+Dox) or treated and withdrawn from doxycycline for 4 days (-Dox 4d). Malignant cells from each tumor type segregated into distinct Clusters consistent with gene signatures for KRAS dependency, cell cycle, or immunomodulation. Numbers on bottom correspond to percent of tumor cell population designated to a specific cluster. UMAP projection images shown below for three marker genes in tumor cells of iKRAS +Dox clusters 1-3.

To extend these observations, we performed single cell RNA sequencing (scRNASeq) of iKRAS control and STAT3 KO orthotopic tumors formed in syngeneic mice administered doxycycline. Tumors were pooled from individual mice and high dimensional sequencing data was embedded using two-dimensional Uniform Manifold Approximation and Projection (UMAP). Unsupervised density clustering segregated cells corresponding to populations of malignant tumor cells, tumor associated macrophages (TAMs), cancer-associated fibroblasts (CAFs), granulocytes/neutrophils, T cells, B cells, endothelial cells, and normal pancreatic cells (Fig. 4D). Tumors derived from doxycycline treated mice bearing STAT3 KO cells contained a relatively lower percentage of malignant tumor cells (20% vs 37%) and a higher percentage of immune cells than tumors from control iKRAS cells (Fig. 4E). Since malignant cancer cells in the tumors dictate growth and context, we evaluated gene expression in the tumor cells. Differential gene expression was performed against aggregated scRNASeq datasets of malignant tumor cells from mice treated with doxycycline or withdrawn from the drug for four days. Evaluation of a subset of immunomodulatory genes showed higher transcript levels in the iKRAS malignant tumor cells relative to STAT3 KO malignant cells, and this was most apparent prior to the removal of doxycycline (e.g. IL10, Arg1) (Fig. 4F). Although both iKRAS and STAT3 KO cells formed tumors when mutant KRAS was turned on, there was an overall trend of higher immune suppression in the iKRAS control cells with functional STAT3. Analysis of infiltrating TAMs and CD8a T cells in the tumors showed modest differences. TAMs in the iKRAS tumors showed a reduction in pro-inflammatory gene expression (M1 polarization) and increase in genes associated with alternative activation (M2 polarization) compared with STAT3 KO tumors (Fig. S3C)(Boutilier & Elsawa, 2021; Sica & Mantovani, 2012). The CD8a T cell population of iKRAS tumors exhibited higher expression of some markers indicative of dysfunction/exhaustion (e.g.Cd160, Lag3, Gmzb) compared to STAT3 KO tumors, and lower expression of some markers for activation (e.g. Il2, Bcl2) (Fig. S3D)(Andreatta *et al*, 2021; Kallies *et al*, 2020). Overall, the data indicate that STAT3 activity contributes to the ability of PDAC tumor cells to evade immunity.

Intratumor heterogeneity is a challenge to successful therapies in human PDAC(Verbeke, 2016). To investigate heterogeneity in the malignant iKRAS tumor cell population, we interrogated the scRNASeq data for gene signatures that correspond to KRAS dependency, cell cycle, and cytokine production (Fig. 4G). Intratumor heterogeneity within the PDAC malignant tumor cell population could be distinguished by patterns of gene expression. The iKRAS tumor cells from doxycycline treated mice displayed three main clusters: Cluster 1, enriched for expression of murine KRAS dependency genes (D’Amico *et al*, 2023; Ischenko *et al*., 2021); Cluster 2, enriched for cell cycle genes; and Cluster 3, enriched for immune modulatory cytokines. Although KRAS signaling is often correlated with proliferation, the analyses indicate these signatures are expressed primarily in distinct tumor cell populations. In addition, the cytokine modulatory cluster does not overlap with either KRAS dependency or cell cycle. It remains to be determined whether there is transition between tumor cell cluster types. The finding of distinct cancer cell clusters highlights the need for combination therapies that target mutant KRAS together with cell proliferation and immune suppression for successful intervention. The STAT3 KO tumor cells lacked the distinct immune cytokine expression of cluster 3. Removal of doxycycline for only four days reduced gene signatures to two clusters for both iKRAS and STAT3 KO cancer cells. The results indicate STAT3 plays a role in immune suppression by the iKRAS malignant tumor cells.

## Discussion tumors

A major question in the aim to cure pancreatic cancer is why loss of oncogenic KRAS does not completely abolish the tumorigenesis of advanced stage cancers. Our findings support an underlying role of STAT3 in maintenance of oncogenic identity following KRAS inhibition in PDAC. The premise of treatment strategies has been that all KRAS-mutant cancers depend on the sustained function of KRAS. However, clinical approaches to target mutant KRAS have not been as successful as hoped and this calls into question the efficacy of single target drugs (Punekar et al., 2022). Our data challenge the paradigm that KRAS is indispensable for tumor maintenance, revealing STAT3 as a compensatory mechanism that sustains malignancy.

Genetic ablation of mutant KRAS was used to mimic pharmacological inactivation as a tool to uncover PDAC tumor vulnerabilities. We previously showed that KRAS depleted PDAC cancer cells have a greatly diminished ability to form tumors in an immunocompetent mouse model (Ischenko *et al*., 2021). In this study we show that KRAS depleted cancer cells that are still tumorigenic depend on the function of STAT3. STAT3 is not an oncogene, however following loss of mutant KRAS, ablation of STAT3 in the murine KPC model abolished the tumorigenic phenotype in immunocompetent mice. Even in human PANC-1 cells that contain a dozen driver mutations (cBioPortal; Cancer Cell Line Encyclopedia), formation of tumors by KRAS depleted cancer cells was dependent on expression of STAT3.

We reveal a role of STAT3 in sustaining tumor cell identity, rather than promoting cell transformation, in KRAS depleted pancreatic cancer cells. Prior studies of PDAC tumor cells depleted for KRAS uncovered transcriptome alterations in key regulatory networks involved in functions of cell fate determination, self-renewal, and differentiation (He *et al*, 2018; Kaestner, 2010; Kinisu *et al*, 2021; Novak *et al*, 2020; Takahashi & Yamanaka, 2006). Previously we described a partial overlap in genes expressed in response to mutant KRAS or STAT3, although STAT3 is not a direct downstream effector of KRAS (D’Amico *et al*., 2023). Here we find that ablation of both mutant KRAS and STAT3 (DKO) disrupts a core program of transcription factors including the pioneer factors FOXA1 and FOXA2. FOXA1/2 are essential for differentiation of endodermal-derived tissues and pancreas development, and they have been reported to contribute to PDAC malignant phenotype (Gao *et al*, 2008; Geusz *et al*, 2021; Kaestner, 2010; Milan *et al*, 2019; Roe *et al*, 2017; Song *et al*, 2010). Most of all, we were able to partially restore tumorigenicity to the DKO cells with transduction of mutant KRAS and STAT3, or by increased expression of FOXA1 and FOXA2. The results demonstrate a plasticity in the tumorigenic properties of the cells, and that loss of KRAS and STAT3 (or their downstream targets) significantly impacts tumorigenic growth. The central finding supports the existence of an embedded transcriptional program of self-renewal that evolved with oncogenic KRAS and is maintained by STAT3 following the loss of KRAS.

Since the DKO KPC cells do not produce tumors in wildtype mice and cannot be analyzed, we evaluated an inducible mutant KRAS model (iKRAS) (Collins *et al*, 2012a; Collins *et al*., 2012b). A limitation of this model is that transient expression of mutant KRAS does not reflect tumor evolution with reduced dependency on KRAS and increased reliance on other pathways. That said, STAT3 expression was also consequential in the iKRAS model. Independent STAT3 KO iKRAS cell clones exhibited reduced tumor forming frequency when transplanted into the pancreas of wildtype mice. When evaluated for short-term regression, the median weight of STAT3 KO iKRAS tumors was reduced compared to controls as early as four days after doxycycline withdrawal. The single cell RNA-seq comparisons of tumors formed by STAT3 KO iKRAS cells with doxycycline induction revealed a relative reduced number of malignant cancer cells in the tumors and an increase in immune cell infiltration. Analysis of the gene signatures for KRAS dependency, cell cycle, or immunomodulation revealed the presence of three distinct clusters of malignant cancer cells in tumors formed by iKRAS cells. The malignant cells from the STAT3 KO tumors lacked an immunomodulatory cytokine cluster, providing further evidence that STAT3 contributes to immune suppression. The heterogeneity found within the malignant cell component of PDAC tumors has been considered a challenge to successful therapeutic intervention (Bailey *et al*, 2016; Collisson *et al*, 2011; Moffitt *et al*, 2015; Peng *et al*., 2019). The distinct malignant cell clusters that we identified by gene signatures in the iKRAS tumors supports the need for combination targeting of KRAS, cell proliferation, and immune evasion.

Suppressing STAT3 function alongside KRAS inhibitors presents a potential therapeutic strategy for overcoming resistance in PDAC. Given the challenge of direct STAT3 inhibition, alternative approaches such as targeting STAT3-regulated genes warrants exploration. Inhibitors of Janus kinases, classical upstream activators of STAT3, have recently been reported to improve efficacy of some anti-cancer immunotherapies (Mathew *et al*, 2024; Zak *et al*, 2024). However, administration of the Janus kinase (JAK) inhibitor tofacitnib did not affect regression of KPC tumors (data not shown). This is not unexpected since JAK inhibitors reduce immune cell function. The limited clinical success of KRAS inhibitors as monotherapies points to the need to identify multiple means by which tumors maintain stemness or immune evasion. We provide evidence for the role of STAT3 in maintaining oncogenic identity and immune suppression following depletion of mutant KRAS.

## Materials & Methods

### Mammalian cells and regents

Murine KRAS^G12D^ p53^R172H^ (KPC) cells (Hingorani *et al*., 2005; Li *et al*., 2018) and inducible KRAS^G12D^ p53^R172H^ (iKRAS) pancreatic epithelial cells (A9993) (Collins *et al*., 2012b) were gifts. Human PANC-1, Phoenix-E and HEK293T cells were obtained from ATCC. All cells were maintained in DMEM supplemented with 5% FBS and 1x antibiotic/antimycotic (Gibco). For proliferation assays, cells were grown for 96 hours, counted with Vi-CELL XR Analyzer (Beckman Coulter), and doubling times were calculated. Cell viability was measured by trypan blue staining. KRAS transgene expression in iKRAS cells was induced by incubating with 1 ug/ml doxycycline hyclate (Sigma) for 24 hours. Stable cell lines were generated and isolated by FACS isolation for GFP or selection in the appropriate antibiotic, puromycin (2 ug/ml) or hygromycin B (100 ug/ml) followed by single clone isolation using glass cloning cylinders.

### Lentiviruses and plasmids

For isogenic CRISPR/Cas9-mediated murine knockout cells, we used murine sgKRAS RNA (5′-gtggttggagctgatggcgt-3′) and sgSTAT3 RNA (5’-gcagctggacacacgctacc-3’ or 5’-gtacagcgacagcttcccca-3’) cloned into LentiCRISPRv2 puro (Addgene)(Mou *et al*, 2017; Sanjana *et al*, 2014). Human knockout cells were generated using sgKRAS RNA (5′-gtagttggagctgatggcgt-3′)(Muzumdar *et al*., 2017) and sgSTAT3 RNA (5’-tgtacagcaccggccgatg -3’). Genomic DNA from putative knockouts was isolated using the Wizard Genomic DNA Purification Kit (Promega), PCR-amplified, cloned into pBluescript KS (+) (Addgene) and confirmed by Sanger sequencing. Primer sequences are available upon request. Retroviral plasmids encoding murine FOXA1, FOXA2, GATA5, HNF1A, HNF4A, KRAS G12D, c-MYC and luciferase were obtained from Addgene. The lentiviral IRES GFP vector (adapted from the pWPXL/pEF1a backbone) encoding murine STAT3 alleles was used as previously described(D’Amico *et al*., 2018). Phoenix-E and HEK293T cells were transiently transfected with the TransIT-LT1 (Mirus) transfection reagent according to manufacturer’s instructions. Production and collection of recombinant retroviruses was done according to standard protocols.

### Histology and immunohistochemistry

Tissues were excised, immersion fixed in ≥5 volumes of 4% paraformaldehyde for 48 hours, transferred to 70% ethanol, and processed by the Stony Brook University Research Histology Core. Paraffin-embedded formalin-fixed 5 μm sections were stained with hematoxylin and eosin (H&E) for histology. Human pancreatic adenocarcinoma samples were obtained as paraffin-embedded tissue specimens from the Stony Brook Medicine BioBank and prepared by the Histology Core. Immunohistochemistry was performed by iHisto (Salem, MA) and Histowiz (Brooklyn, NY). Slides were scored independently by two investigators. H-scores represent the sum of malignant cell staining intensity in the tumors (negative 0, weak 1, moderate 2 or strong 3) multiplied by the percentage of stained cells.

### Expression analyses

Western blotting was performed using whole cell extracts prepared by lysing cells in buffer containing 10 mM Tris-HCl, pH7.4, 150 mM NaCl, 1 mM EDTA, 10% glycerol, 1% Triton X100, 40 mM NaVO4, 0.1% SDS, and 1x protease inhibitors (Roche). Nuclear and cytoplasmic fractions were prepared using NE-PER nuclear and cytoplasmic extraction reagents (ThermoFisher). Proteins were detected with antibodies to P-ERK1/2 (9101), E-cadherin (3195), c-FOS (2250), FOXA2 (8186), KLF4 (4038), KLF5 (51586), P-MEK1/2 S217/221 (9154), MYC (5605), NFkB p65 (8242), RASG12D (14429), α-SMA (19245), STAT3 (9139), P-STAT3 Y705 (9145) (Cell Signaling); KRAS (sc-30, Santa Cruz); HNF4A (A11496, ABclonal); GATA5 (R12-2154, AssayBioTech); ERK1/2 (05-157, EMD Millipore); FOXA1 (ab170933, Abcam). Total cellular RNA was isolated using PureLink RNA (ThermoFisher) according to manufacturer’s specifications and phenol-chloroform extracted. Pancreatic tumor tissue was incubated overnight at 4°C in ≥5 volumes of RNA*later* solution (ThermoFisher). Whole transcriptome RNA-sequencing (RNA-seq) with bioinformatics was performed by Novogene Corp. (http://en.novogene.com). Gene ontology (GO) classifications were generated by Novogene or the online GO Enrichment Analysis tool (http://geneontology.org).

Human pancreatic adenocarcinoma clinical data were downloaded from cBioPortal (http://www.cbioportal.org) or from the Amsterdam UMC (AUMC) database (Dijk et al). Tumor samples were classified as KRAS dependent/KRAS-type, or KRAS independent/RSK-type by calculating the sum of RNA expression values (z-scores) using previously characterized gene modules (Singh *et al*., 2009; Yuan *et al*., 2018). A murine KRAS dependency signature was used to calculate RAS dependency scores and was previously described (D’Amico *et al*., 2024; Ischenko *et al*., 2021). STAT3 signature scores were computed from a set of regulated genes as reported(Dauer *et al*., 2005). Additional pathway activity scores were determined using curated hallmark gene sets from the Molecular Signatures Database (www.gsea-msigdb.org) based on the average fpkm values (stem module M1999; hallmark EMT M5930, and cell adhesion M9500). Gene Set enrichment analysis was performed using the application available from the Broad Institute (http://software.broadinstitute.org/gsea/. Scatter plots and heatmaps were generated using online Heatmapper software or Microsoft Excel.

### Tumorigenicity in mice

All animal studies complied with ethical regulations for animal testing and research approved by the Institutional Animal Care and Use Committee of Stony Brook University (protocol 2011-0356). We used adult (8-10 wks old) male and female mice from The Jackson Laboratory, strains C57BL/6J, FVB/NJ, and NU/J. Subcutaneous implantations were performed using 10^4^ murine or 4 x 10^5^ human cells in 100 μl of Matrigel diluted 1:10 with Opti-MEM (Corning). Tumor latency was defined as the period between implantation of tumorigenic cells and the appearance of tumors ∼1mm^3^. Orthotopic injections into the pancreas were performed with 10^4^ cells in diluted Matrigel using standard procedures (Kim *et al*, 2009). The animals were observed for tumor development by palpation or bioluminescence. The frequency of tumor-initiating cells was calculated using online ELDA software (http://bioinf.wehi.edu.au/software/elda/), and the number of cells implanted subcutaneously ranged from 10^2^ to 10^4^ (Hu & Smyth, 2009). For iKRAS tumor formation, the drinking water was provided *ad libitum* and supplemented with 2% sucrose, doxycycline hyclate (0.5 mg/ml), neomycin (0.5 mg/mL) and ampicillin (1 mg/mL). Water was changed every 3-4 days. For bioluminescence imaging, mice were injected intraperitoneally with 200 μl 15 mg/ml D-luciferin (Syd Lab Inc.) and imaged using the Lumina III in vivo imaging system (IVIS) with Living Image Software (PerkinElmer). Drug treatment cocktails were administered by IP injection and were composed of anti-PD1 (BE0146), anti-CTLA4 (BE0164), and anti-CD40 (BE0016) antibodies from Bio X Cell at 1 mg/kg in PBS, and with MRTX1133 (Chemietek) at 10 mg/kg and GSK1120212 (Selleck Chem llc) at 2 mg/kg in 5% DMSO, 40% PEG300, 5% Tween-80, and 45% saline. Controls received rat IgG2a antibodies (Bio X Cell). For flow cytometry, tumor cells were dissociated with 1X collagenase/hyaluronidase (Stem Cell Technologies). Cells were incubated with unconjugated anti-CD16/CD32 prior to staining with the following fluorophore-conjugated antibodies from BioLegend, Invitrogen, or BD Biosciences: anti-CD45 (APC-eF780), anti-TCRβ (PerCp/Cy5.5), anti-CD8α (BV786), anti-CD4 (BV605), anti-CD3e (BV421), anti-TCRd (PE), anti-CD11b (APC), anti-F4/80 (PE/Cy7) and anti-Ly6G (PE/Dazzle594). Live/dead dye was purchased from Invitrogen. Cells were stained protected from light at 4°C for 20 minutes. After staining, cells were fixed with 2% paraformaldehyde (Electron Microscopy Sciences) for 20 minutes prior to data acquisition on a LSRFortessa (BD) or an Aurora (Cytek). Data were analyzed with FlowJo software (Tree Star). Data represent three experiments with tumors pooled from 2-3 mice.

### scRNA-Seq

Orthotopic tumors formed in FVB male mice with iKRAS parental control (A9993)(Collins *et al*., 2012b) or iKRAS STAT3 KO cells were pooled from 2-3 individual mice at day 0 (∼300-400mg) or following doxycycline removal for 4 days (day 4) (∼100-300mg). Single cell suspensions were prepared by finely mincing tumor tissue with a razor blade followed by incubation with 1x collagenase/hyaluronidase (StemCell Technology), 0.5 mg/ml Liberase-DL (Roche), and 0.1 mg/ml DNase I (Sigma) in DMEM for 45 minutes at 37°C with continuous mixing. Single-cell RNA sequencing and analysis was performed by the Cold Spring Harbor Laboratory Single-Cell Biology Shared Resource, Genomics Technology Development. Cells were loaded into the 10x Genomics microfluidics device along with 10X Genomics gel beads (kit v2 PN-120237) containing barcoded oligonucleotides, reverse transcription (RT) reagents, and oil, resulting in gel beads in emulsion. The scRNA-seq library preparation followed the manufacturer’s protocol (10x Genomics) using the Chromium Single Cell 3-Library. Libraries were paired end sequenced using the Illumina HiSeq 4000 sequencing system. Genomics Cell Ranger pipeline (version 3.0.1) was used to convert single-cell sequencing files and secondary analysis was with Loupe Cell Browser v3.0.1 (10X Genomics).

## Statistics and reproducibility

Statistical analysis was performed using two-tailed Student’s t-test, Fisher’s exact test, or Wilcoxon test, as appropriate for the dataset. An FDR-adjusted p-value (q-value) was calculated for multiple comparison correction. Individual mice, tumors, and tumor cell lines were considered biological replicates. Technical replicates were in triplicate. Statistical details for each experiment are denoted in the corresponding figures and figure legends. The micrographs (H&E and IHC images) represent two or more independent experiments. For the quantification of IHC, cells were counted manually with an average of 10-100 fields per tumor. All data are presented as mean ± SD.

## Data availability

Human PDAC expression profiles from The Cancer Genome Atlas (TCGA) were downloaded as z-scores relative to diploid samples from cBioPortal (http://www.cbioportal.org) along with additional tumor and clinical annotations. PDAC datasets from the Amsterdam UMC (AUMC) (Dijk et al), the PanCuRx Translational Research Initiative (COMPASS)(Chan-Seng-Yue *et al*, 2020), and The NCI Clinical Proteomic Tumor Analysis Consortium (CPTAC) were used as described. Data deposited in DRYAD as curated Microsoft Excel files for murine KPC cell RNA-seq data, iKRAS scRNA-seq data, and human PANC-1 RNA-seq data at https://doi.org/doi:10.5061/dryad.1vhhmgqzc, https://datadryad.org/stash/share/Q7KVMuOcvfT70YvD7zd78MOI4nLAj3cMyW_Lq_dKhoo, and https://doi.org/10.5061/dryad.5hqbzkhct. The scRNA-seq data have been deposited in the GEO database under accession code GSE275858. Additional information and/or reagents are available from the authors on request.

## Acknowledgements

This work was supported by NIH grant RO1CA236389, the Carol M. Baldwin Breast Cancer Research Award, and Stony Brook Center for Healthy Aging Award to NCR. We wish to acknowledge Dr. Oleksi Petrenko for his help in data analysis and advice with the investigation and manuscript. We also wish to thank Dr. Jinyu Li for assistance with data analysis, Jean Rooney in the Stony Brook University Division of Laboratory Animal Research for technical assistance in mouse surgeries, Yan Ji and technical support provided by the Research Histology Core Laboratory in the Department of Pathology, Dr. Richard Kew, Scientific Director of Stony Brook Medicine’s Biobank in the Department of Pathology and Cancer Center, and Dr. Jonathan Preall, Head of Genomics Technology Development Cold Spring Harbor Laboratory (CSHL) for single-cell RNA-seq and bioinformatics analyses. We very much appreciate the generosity of Dr. David Tuveson (CSHL) and Dr. Ben Stanger (University of Pennsylvannia) for providing KPC cells lines, and Dr. Marina Pasca di Magliano (University of Michigan) for providing iKRAS cell lines.

## Supplemental Figure Legends

**Supplemental Figure 1.**
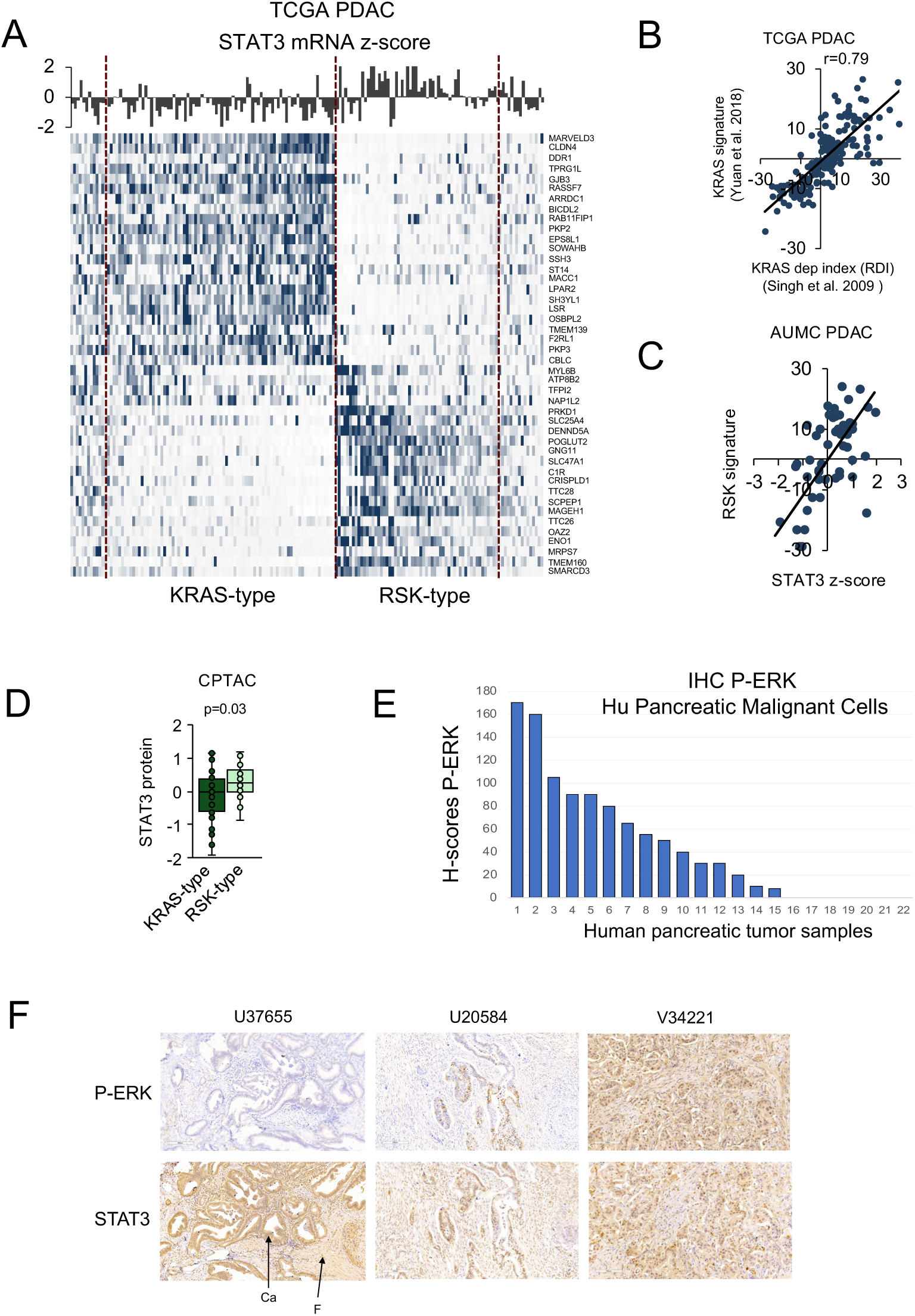
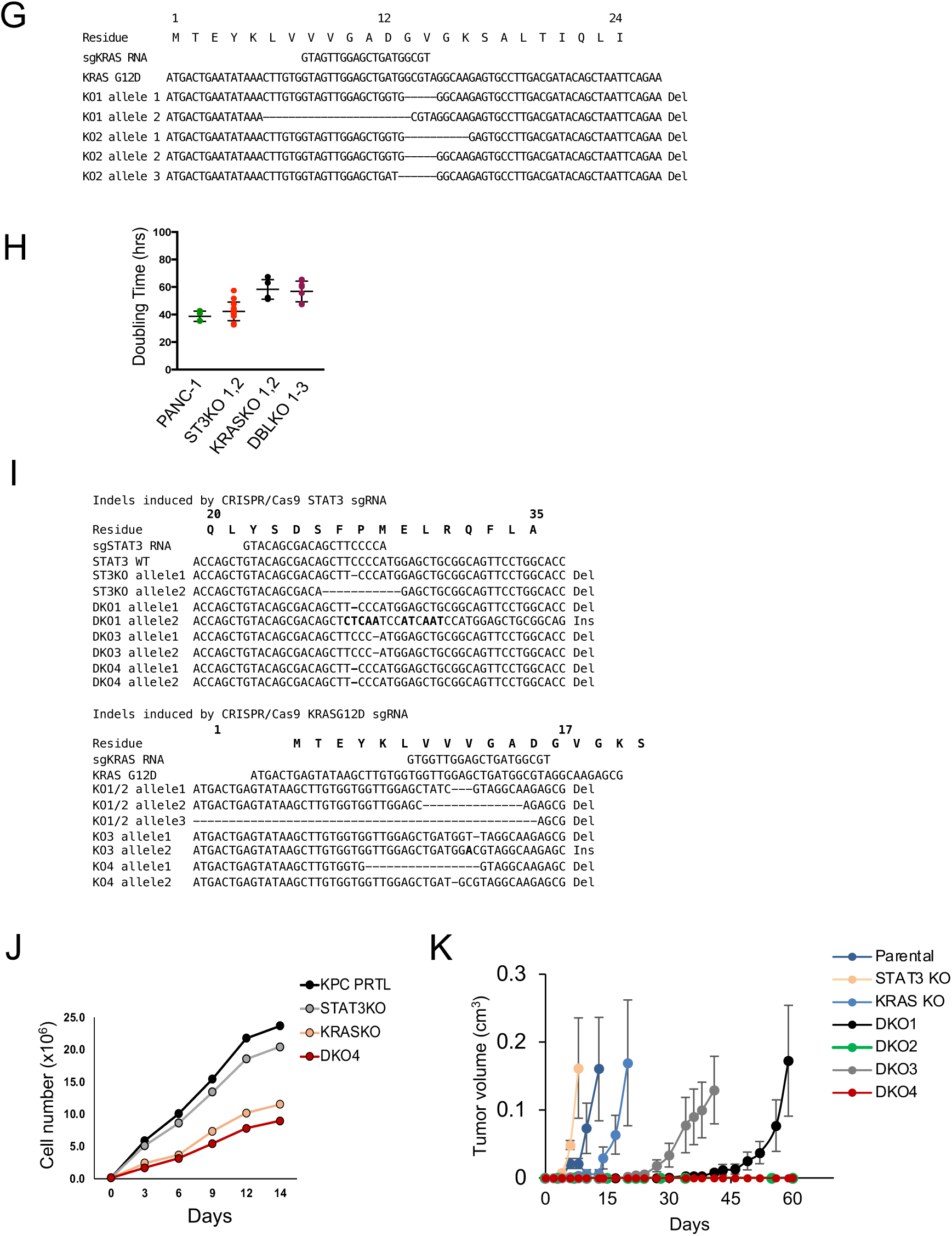
STAT3 supports tumorigenicity of mutant KRAS depleted PDAC. S1A. Gene expression profiling of 168 human pancreatic adenocarcinomas in the TCGA PanCancer Atlas database corresponding to a gene signature of KRAS dependency (KRAS-type) or reduced KRAS dependency (RSK-type)(Yuan *et al*., 2018). STAT3 mRNA expression z-score for each tumor sample is displayed on top. S1B. Scatter plot showing correlation of two derived KRAS dependency gene signatures with references for 150 human pancreatic adenocarcinomas in the TCGA database. S1C. Scatter plot showing correlation of STAT3 mRNA expression and RSK-type gene signature (Yuan *et al*., 2018) in human pancreatic ductal adenocarcinomas from the AUMC (Amsterdam University Medical Centers) database (n=80). S1D. Proteomic data from human pancreatic adenocarcinomas of the NCI Clinical Proteomic Tumor Analysis Consortium (CPTAC) comparing levels of STAT3 protein in TCGA tumors classified as KRAS-type or RSK-type (n=40). S1E. Histo-score (H-score) of the cancer cells in 22 human pancreatic adenocarcinoma tumors from the Stony Brook Medicine Biobank following immunohistochemistry staining for phosphorylated-ERK. S1F. Representative images of IHC staining of human pancreatic tumors scored in Fig. S1E with antibodies to phospho-ERK (P-ERK) or STAT3. Tumor cancer cells (Ca) and fibroblast stromal cells (F) are indicated in the tumor. Scale bar 100mm. S1G. Representative sequences of major deletions created by CRISPR-mediated editing of mutant KRAS in the human PANC-1 cell line, determined following PCR amplification and cloning. S1H. Growth in culture of PANC-1 parental cells expressing empty vector or knockout derivatives shown. Graph displays doubling time. S1I. Representative sequences of the major indels modified by CRISPR-mediated gene editing of STAT3 and mutant KRAS in the KPC cell lines were determined following PCR amplification and cloning. S1J. Growth in culture of KPC parental cells (prtl) and derived STAT3 KO, KRAS KO, and DKO cl4 cells. Cell number during course of two weeks is shown. S1K. Relative tumor development during 60 days of parental KPC cells and derived STAT3 KO, KRAS KO and four DKO KPC cell lines following subcutaneous implantation in nude mice (n>8).

**Supplemental Figure 2.**
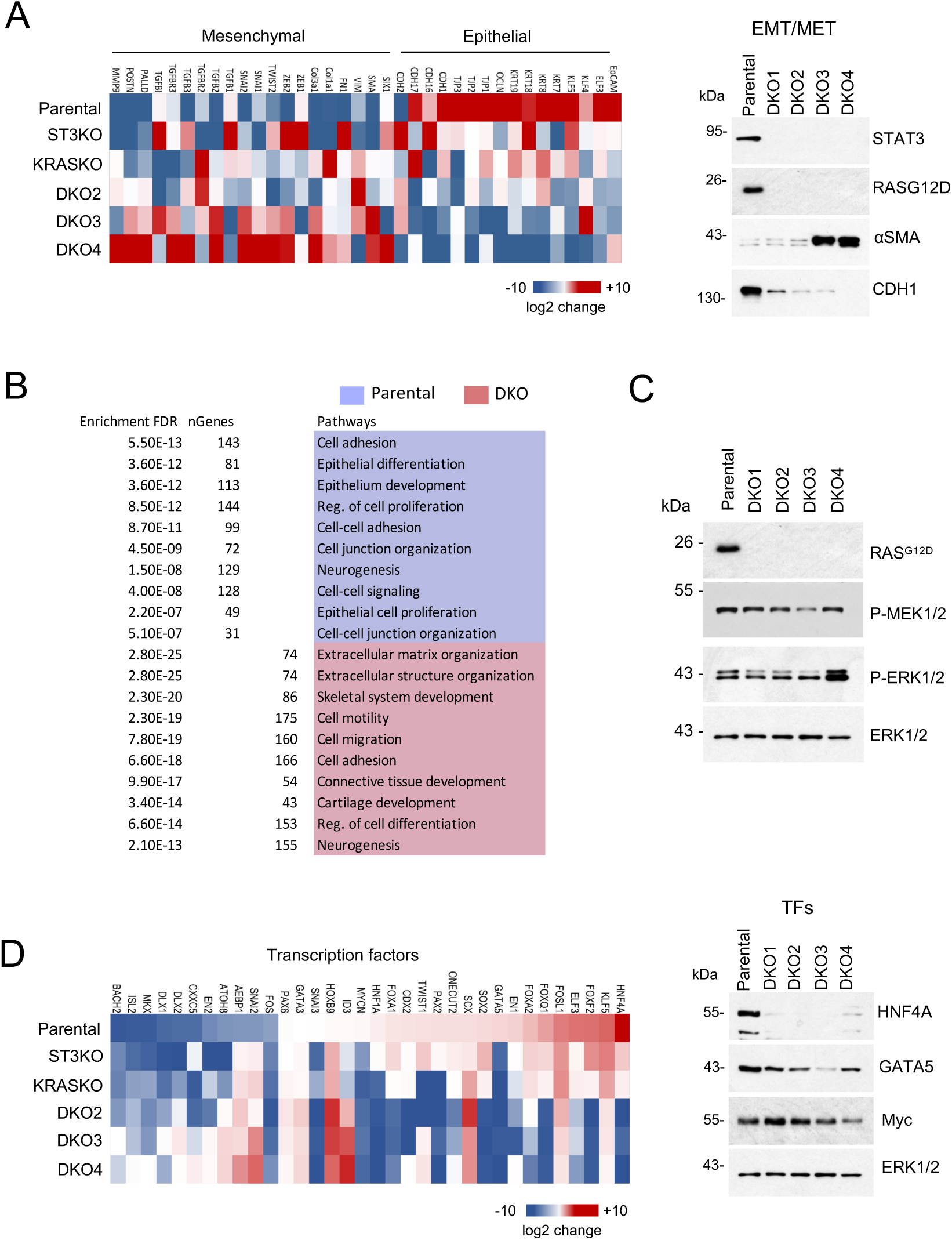
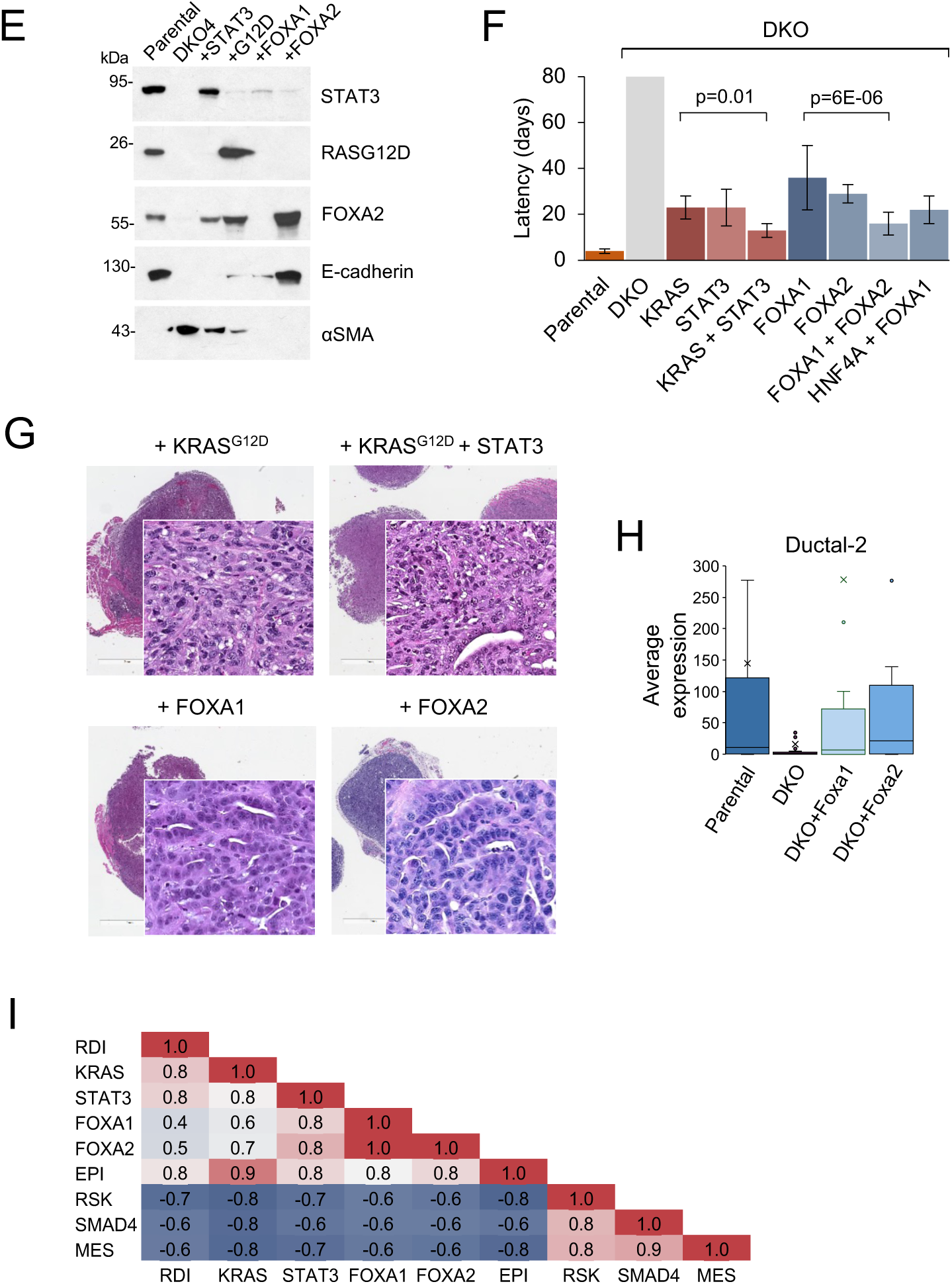
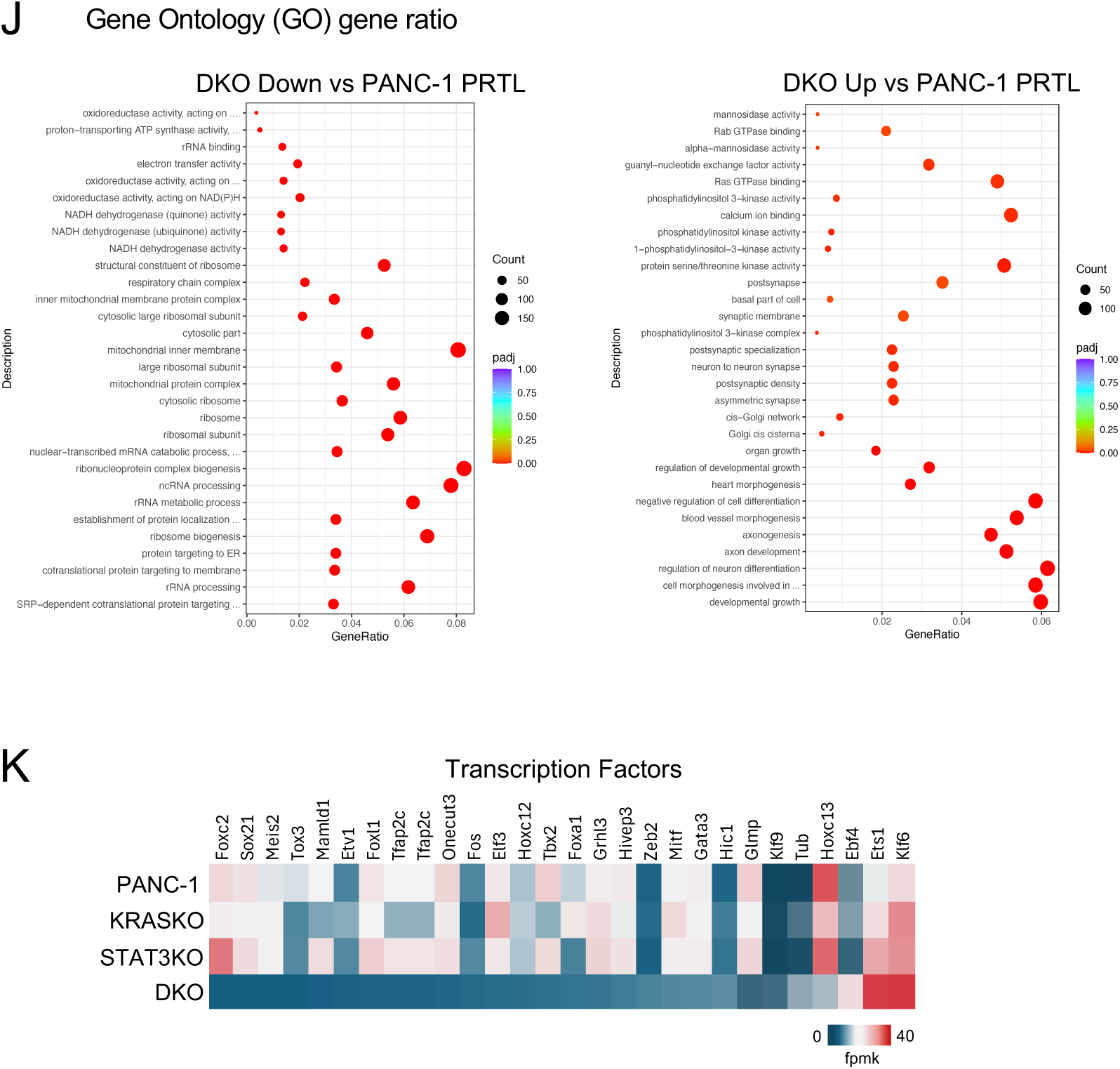
Transcriptional reprogramming following loss of KRAS and STAT3. S2A. Left) Heatmap of gene signatures for mesenchymal and epithelial identity derived from RNA-seq data of KPC parental, STAT3 KO cells, KRAS KO cells, and three of the DKO clones. Right) Western blots of KPC parental cells and four DKO cell lines confirming RNA-seq data for expression of epithelial E-cadherin (CDH1) or mesenchymal smooth muscle actin (αSMA). S2B. Gene ontology (GO) classification of top biological processes that are upregulated in KPC parental cells or DKO4 cells. S2C. Western blot of KPC parental cells and four DKO derived clones for phosphorylation activity of MEK and ERK1/2. S2D. Left) Heatmaps of differential gene expression from RNA Seq analyses corresponding to a set of transcription factors in parental KPC cells, STAT3 KO cells, KRAS KO and three DKO derived clones. Right) Western blots of KPC parental cells and four DKO cell lines for HNF4A, GATA5, and MYC. ERK1/2 are used as loading controls. S2E. Representative Western blot of parental KPC cells, DKO cells, and DKO cells restored for expression of the designated gene by lentiviral transduction. S2F. Restoration of tumorigenicity in nude mice of DKO4 cells quantified by latency in days (∼1mm tumor) following transduction with mutant KRAS, STAT3, FOXA1, FOXA2 or the indicated combinations compared with parental KPC cells. S2G. Representative H&E histology images of tumors formed following transduction of DKO cells with genes noted. S2H. Average expression of the top ductal-2 type PDAC genes determined by RNA-seq of KPC parental cells, DKO cells, or DKO cells transduced with Foxa1 or Foxa2 (Peng *et al*., 2019). S2I. Pearson’s coefficient analyses aligning gene expression signatures of reconstituted KPC cells as noted with PDAC TCGA tumor samples (n=168, stage I/II tumors) according to RAS dependency (RDI), molecular subtype (i.e. KRAS-type vs RSK-type), and epithelial (EPI) or mesenchymal (MES) status. S2J. Gene ontology (GO) comparison of biological processes expressed in human PANC-1 DKO cells versus parental PANC-1 cells derived from RNASeq analyses. Comparisons shown as dot matrices. S2K. Heatmaps of differential gene expression from RNA Seq analyses corresponding to a set of transcription factors in parental human PANC-1 cells or derived KRAS KO, STAT3 KO, or DKO cells.

**Supplemental Figure 3.**
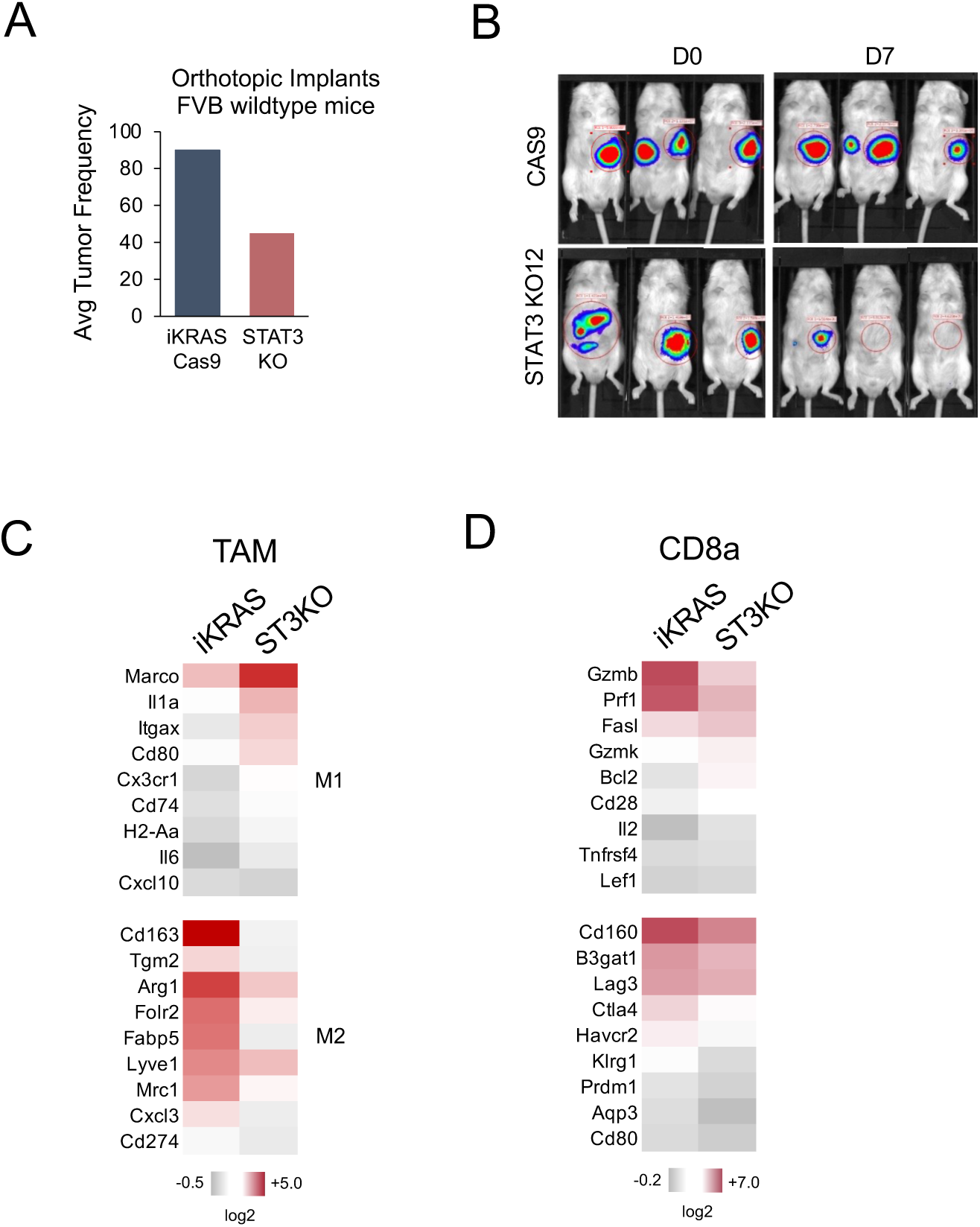
STAT3 depletion in an Inducible mutant KRAS PDAC model. S3A. Average frequency of tumor formation in FVB wildtype mice treated with doxycycline of parental iKRAS cells expressing empty vector (EV) or three independent CRISPR-edited STAT3 KO iKRAS clones. S3B. Representative *in vivo* bioluminescence imaging of tumors formed by orthotopic implants of iKRAS parental cells expressing empty vector (Cas9) or derived STAT3 KO12 cells that were transduced to express luciferase. Mice were administered doxycycline and tumors were evaluated on day 0 (D0) or seven days after withdrawal of doxycycline (D7). S3C. Comparative heatmaps of scRNA-seq individual datasets for a subset of genes expressed in the tumor associated macrophages (TAMs) in tumors formed by iKRAS control cells or STAT3KO derived cells in FVB mice treated with doxycycline. S3D. Comparative heatmaps of scRNA-seq individual datasets for a subset of genes expressed in CD8a+ T cells (CD8a expression log2 fold greater than CD4 expression) in tumors formed by iKRAS control cells or STAT3KO derived cells in FVB mice treated with doxycycline.

## Notes

### Competing Interest Statement

The authors have declared no competing interest.

